# Activation of goblet cell stress sensor IRE1β is controlled by the mucin chaperone AGR2

**DOI:** 10.1101/2023.07.06.547951

**Authors:** Eva Cloots, Phaedra Guilbert, Mathias Provost, Lisa Neidhardt, Evelien Van de Velde, Farzaneh Fayazpour, Delphine De Sutter, Savvas N. Savvides, Sven Eyckerman, Sophie Janssens

## Abstract

As secretory cells specialized in the production of mucins, intestinal goblet cells are challenged by the need for efficient protein folding. Goblet cells express Inositol-Requiring Enzyme 1β (IRE1β), a unique unfolded protein response (UPR) sensor that is part of an adaptive mechanism that regulates the demands of mucin production and secretion. However, how IRE1β activity is tuned to mucus folding load remains unknown. We identified the disulfide isomerase and mucin chaperone AGR2 as a goblet cell specific protein that crucially regulates IRE1β-, but not IRE1α-mediated signaling. AGR2 binding to IRE1β disrupts IRE1β dimerization, thereby blocking its downstream endonuclease activity. Depletion of endogenous AGR2 from goblet cells induces spontaneous IRE1β activation, suggesting that alterations in AGR2 availability in the endoplasmic reticulum sets the threshold for IRE1β activation. We found that AGR2 mutants lacking their catalytic cysteine or displaying the disease-associated mutation H117Y were no longer able to dampen IRE1β activity. Collectively, these results demonstrate that AGR2 is a central chaperone regulating the goblet cell UPR by acting as a rheostat of IRE1β endonuclease activity.

## Introduction

The gastrointestinal (GI) tract is protected by an essential mucus layer, which forms the main physical barrier between the intestinal epithelium and the outside world. The main protein component of the intestinal mucus layer is the glycoprotein MUC2, which is produced by goblet cells, specialized secretory cells that are present both in the small and large intestine ^1^. MUC2 is a large carbohydrate of over 5100 amino acids that becomes assembled in disulfide bond stabilized dimers in the endoplasmic reticulum (ER) before translocating to the Golgi ^1–3^. As such, its folding poses a significant burden on the ER, making goblet cells exquisitely sensitive to folding defects ^4^. This is further supported by the close link between aberrations in mucus folding capacity and inflammatory bowel disease ^4–6^.

To ensure proper folding, eukaryotes developed a sophisticated adaptive response termed the unfolded protein response (UPR) orchestrated by three ER-based sensors that respond to changes in folding load: PKR-like ER Kinase (PERK), Activating Transcription factor (ATF)6 and inositol requiring enzyme-1 (IRE1)(gene name *ERN1*) ^7^. Together, they coordinate a program to slow translation, expand the ER, upregulate the expression of chaperones, and amplify the capacity to process misfolded proteins for degradation. IRE1 is the most evolutionarily conserved member of the three UPR sensors and is characterized by an N-terminal ER-luminal sensor domain, a Type I transmembrane domain, and two C-terminal cytoplasmic enzymatic domains: a kinase and an endonuclease domain ^8^. In steady state conditions, IRE1 is kept in an inactive monomeric state by binding the chaperone Binding Immunoglobulin Protein (BIP) ^9^. Upon accumulation of unfolded proteins in the ER, BIP dissociates from IRE1α, leading to its default dimerization/oligomerization and transphosphorylation ^10–12^. Nucleotide binding stabilizes the back-to-back dimer interface necessary for endonuclease activity ^13^. While the details have not been fully elucidated yet, binding of unfolded proteins to the luminal domain has been postulated to induce IRE1 oligomerization ^14,15^. This creates a composite RNA binding pocket that accommodates IRE1’s main endonuclease substrate, X-box binding protein 1 (*Xbp1)* mRNA, allowing excision of a 26nt intron. The resulting frameshift leads to the production of the active transcription factor XBP1s ^16^. Apart from *Xbp1*, IRE1α has also been shown to target additional mRNAs for degradation in a process termed Regulated IRE1 Dependent Decay or RIDD, which further relieves the folding burden on the ER ^17,18^.

While IRE1α is ubiquitously expressed, goblet cells and other epithelial cells in the gastrointestinal and respiratory tract, uniquely express a second paralogue of IRE1, IRE1β (gene name *ERN2*) ^8,19,20^, with expression levels that largely exceed those of IRE1α ^21^. IRE1β arose through whole genome duplication in ancestral vertebrates and analysis of IRE1β sequence variations in different vertebrate species suggested that IRE1β adopted a neofunctionalization with the emergence of a mucus-based system of barrier immunity ^8^. This, together with its unique expression in epithelial cells lining mucosal interfaces suggests that IRE1β functions in mucosal homeostasis, which is supported by earlier studies. IRE1β^-/-^ mice are more susceptible to dextran sulphate sodium (DSS)-induced colitis, exhibiting worse inflammation and premature lethality ^22^. Absence of IRE1β leads to a defect in goblet cell numbers ^23^ and a recent study assigned a role for IRE1β in goblet cell maturation, mucin secretion and mucus barrier assembly, that was strictly dependent on the present of an intact microbiome ^24^. At a mechanistic level, Tsuru *et al.* showed that IRE1β is responsible for trimming *Muc2* mRNA pools in a RIDD dependent manner, thereby avoiding overloading of the ER with MUC2 polypeptide chains ^25^. IRE1α and IRE1β show divergent roles in intestinal homeostasis. While loss of IRE1β affects goblet cell number, this has not been observed in IRE1α deficient mice ^23,25^. Furthermore, IRE1β appears to protect from Crohn’s-disease like ileitis, triggered upon absence of XBP1 and ATG16L1 in Paneth cells, while IRE1α contributes to the disease ^23^.

How IRE1β exerts these goblet cell-specific functions remains largely enigmatic. Despite its close homology to IRE1α, IRE1β displays only weak endonuclease activity, and even acts as a negative regulator of IRE1α endonuclease activity, both in vitro ^21^ and in vivo ^23^. IRE1β only marginally responds to classical ER triggers and predominantly forms dimers instead of oligomers, consistent with a presumed prominent role for IRE1β in RIDD rather than in XBP1 splicing ^21,26,27^. One of the most remarkable properties of IRE1β, - and explaining at least partially its poorly understood function -, is the fact that overexpression of the protein in cell culture leads to a hyperactive and deleterious state. HeLa cells undergo rapid cell death upon IRE1β expression, suggested to be caused by 28S rRNA degradation ^28^. At present, it is incompletely understood how IRE1β is regulated and how the difference in activity levels in distinct model systems can be explained.

In this work, we identified the chaperone Anterior-gradient protein homolog 2 (AGR2) as an interactor and essential regulator of IRE1β activity. AGR2 is a protein disulfide isomerase (PDI) uniquely expressed in goblet cells and involved in mucus maturation ^29^. Perturbations in AGR2 cause intestinal pathology due to disruptions in mucin maturation both in murine models ^29^ and in humans ^30^. Here we describe AGR2’s novel role as a regulator of IRE1β activity by keeping the protein in an inactive (monomeric) form, which is conceptually similar as to how the chaperone BIP tunes IRE1α activity. In non-goblet cells, deficient for endogenous AGR2, overexpression of IRE1β leads to spontaneous dimerization and unrestrained activation, causing cell death, which can be rescued by co-expression with exogenous AGR2.

## Results

### IRE1β activity is attenuated upon overexpression in goblet cells

Previous studies on the endonuclease activity of IRE1β, have generally relied on cell types that do not represent the natural environment for IRE1β. Hela cells, where rapid overactivation and even cell death is reported upon exogenous expression of IRE1β ^28^, express only background levels of IRE1β transcript (**Fig 1A**). We therefore selected two cell lines with highly diverging endogenous expression levels of IRE1β at transcript and protein levels, to compare IRE1β endonuclease activity and toxicity upon overexpression. Calu-1 cells are lung epithelial cells that do not express IRE1β at endogenous level, like most other cell lines, whereas the goblet cell-like LS174T line does show endogenous expression of IRE1β transcript and protein (**Fig 1A** and **Fig S1A**). To assess the endonuclease activity of IRE1β specifically without any confounding effects of IRE1α endonuclease activity, both cell lines were engineered by CRISPR-Cas9 for IRE1α deficiency and transduced with a lentiviral, doxycycline-inducible IRE1β expression module (Calu-1^ERN^^1-/-IRE1βFLAG-DOX^ and LS174T^ERN^^1-/-IRE1βFLAG-DOX^) (**Fig 1B**). As reported previously for HeLa cells, overexpression of IRE1β in Calu-1 cells induced increasing cell death over time (**Fig 1C**, top panels at 72h post doxycycline induction). This process could be reversed by adding the IRE1 specific endonuclease inhibitor 4µ8C ^31^, but occurred independently of IRE1α, as both *ERN1*+/+ and *ERN1*-/- Calu-1 cells exhibited IRE1β-mediated toxicity that could be rescued by 4µ8C (**Fig S1B**). All further experiments were performed with *ERN1*-/- cell lines. In contrast to our observations in the lung epithelial Calu-1 cells, overexpression of IRE1β in colon epithelial LS174T cells did not cause cellular toxicity (**Fig 1C**, lower panels), demonstrating that the cellular context strongly influences IRE1β behavior. The difference in phenotype was not caused by a lack of transgene expression in LS174T cells, as an anti-FLAG IRE1β western blot demonstrated even higher transgene expression levels in LS174T cells compared to Calu-1 cells (**Fig 1D)**, which was consistent over time (**Fig S1C**). Notably, in presence of 4µ8C, Calu-1 cells expressing high levels of IRE1β could be readily retrieved (**Fig S1C)**.

**Figure 1.**
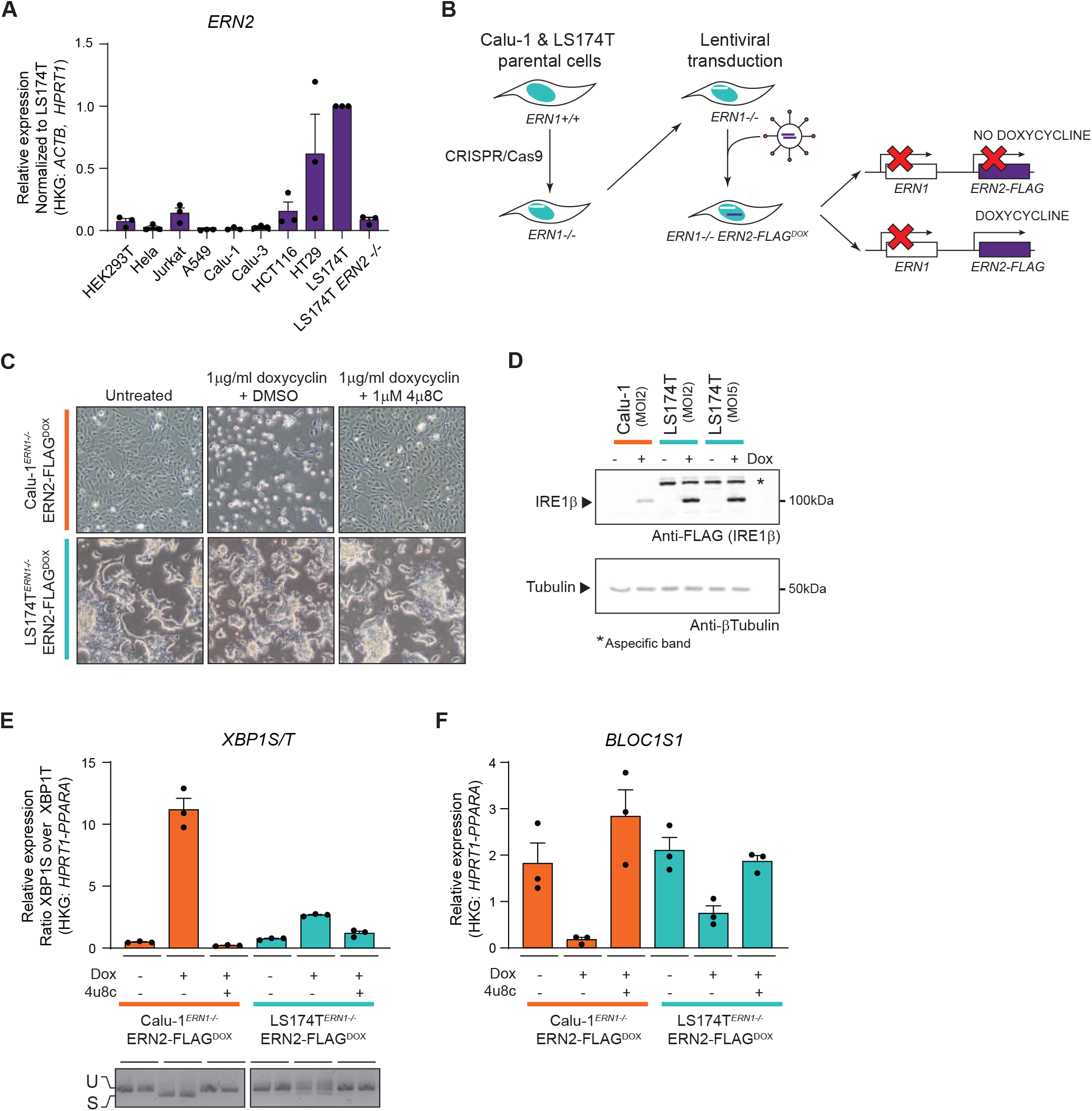
IRE1β activity is attenuated upon overexpression in goblet cells. **A** RT-qPCR analysis of *ERN2* (IRE1β) transcript expression in human cell lines. Cultures were sampled three times and *ERN2* expression is shown relative to the expression in LS174T parental cells. **B** Schematic representation of cell lines used for further studies. First, *ERN1*-/- clones of the LS174T and Calu-1 parental lines were established by CRISPR/Cas9. Then, *ERN1*-/- cells were transduced with a TetOn module for doxycycline controlled IRE1β-FLAG expression. **C** Photographs showing the phenotype of cultures overexpressing IRE1β-FLAG. Left panels show untreated cultures, middle panels show cultures that received 1μg/ml doxycycline for 72 hours (and thus express IRE1β-FLAG), cultures in the right panels received 1μg/ml doxycycline and 1μM IRE1 endonuclease inhibitor 4μ8C. Representative of three independent experiments. **D** Western blot verifying transgene expression in the cell lines used in (C). All remaining adherent cells after 24 hours of transgene induction were collected and lysates were probed for FLAG-IRE1β expression via immunoblot. Tubulin was used as a loading control. Multiplicity of Infection (MOI) indicates the theoretical number of viral particles added. **E, F** RT-qPCR analysis of *XBP1*S/T **(E)** and *BLOC1S1* transcript levels **(F)** after 24 hours of transgene induction. Representative of three independent experiments with three replicates per condition. **(E)** bottom picture shows XBP1 splicing in the same samples assayed by conventional PCR. The fast-migrating band is the spliced XBP1 transcript, and the slow migrating band the XBP1 unspliced transcript.

To measure spontaneous IRE1β endonuclease activity more precisely, XBP1 splicing was quantified in cultures overexpressing IRE1β for 24h. Also in this case, LS174T cells exhibited a severely diminished activity of IRE1β, with Calu-1 cells reaching much higher levels of XBP1 splicing according to qPCR and PCR analyses (**Fig 1E**). Finally, RIDD activity monitored via the degradation of the canonical RIDD target *Bloc1s1* was less induced upon overexpression of IRE1β in LS174T cells compared to Calu-1 cells, albeit not fully disrupted (**Fig 1F**). Overall, these data demonstrate that upon overexpression, IRE1β displays basal activity that is repressed in the context of a goblet cell-like cellular background.

### The mucin chaperone AGR2 is a goblet cell-specific interactor of IRE1β

Higher XBP1 splicing levels in Calu-1 cells compared to LS174T cells could be explained by a regulating factor present in the latter (which, like goblet cells normally express IRE1β), but absent in cell types like Calu-1 (that do not normally express IRE1β). To identify potential candidate proteins that could fulfill this role, affinity-purification mass spectrometry (AP-MS) was performed in LS174T cells using both IRE1β-FLAG and IRE1α-FLAG expressed from the doxycycline inducible system (**Fig 2A**). Co-purified proteins were digested and analyzed by mass spectrometry. Enrichment of peptides in IRE1-FLAG samples compared to empty vector control was assessed by label free quantification (LFQ) and two-sample t-test. After analysis, 26 proteins had a log_2_ fold change (log_2_FC) enrichment over control cells of >2, and log_10_Adj p-val of >2 in the IRE1α-FLAG pulldown samples compared to control cells (**Fig 2B-C**, left panel). For IRE1β-FLAG, 55 proteins remained using the same criteria (**Fig 2B-C**, right panel). In total, 11 proteins were commonly identified for both IRE1 paralogues (**Fig 2B**, purple intersection) including BIP (*HSPA5*), the SEC61 translocon and CDC37, a kinase chaperone previously identified to interact with IRE1α (Mandal et al 2007 JCB, Ota & Wang 2012 J Biol Chem). IRE1β co-precipitated with many ribosomal proteins (in line with IRE1β’s previously reported ability to cleave ribosomal RNA ^28^) and notably, the goblet cell specific protein disulfide isomerase AGR2 (**Fig 2B, light blue region**). Of all the interaction partners uniquely identified for IRE1β, the chaperone AGR2 stood out as AGR2 had been described as an essential mediator of MUC2 processing and secretion ^29^, while IRE1β has been shown to degrade *Muc2* mRNA pools ^25^. Furthermore, their interaction yielded one of the highest fold changes at a log_2_FC of ∼6.5 (see **Fig 2C** right panel and **Supplementary table 1**). For further validation of the specificity of the interaction between IRE1β and AGR2, an antibody-free system was utilized, in which both IRE1 paralogues were biotin-tagged using the Avi-tag ^32^, which can only be biotinylated and pulled down by streptavidin beads upon co-expression of the *E. coli* biotin ligase BirA. Also in this overexpression system in HEK293T cells, AGR2 interacted specifically with IRE1β, while only a very faint signal interacting with IRE1α was retrieved (**Fig 2D**). Finally, the interaction between endogenous AGR2 and IRE1β was confirmed in colon tissue, using *Agr2* deficient mice as a negative control (**Fig 2E**). In summary, these data corroborate the PDI AGR2 as a specific interactor of IRE1β, not IRE1α.

**Figure 2.**
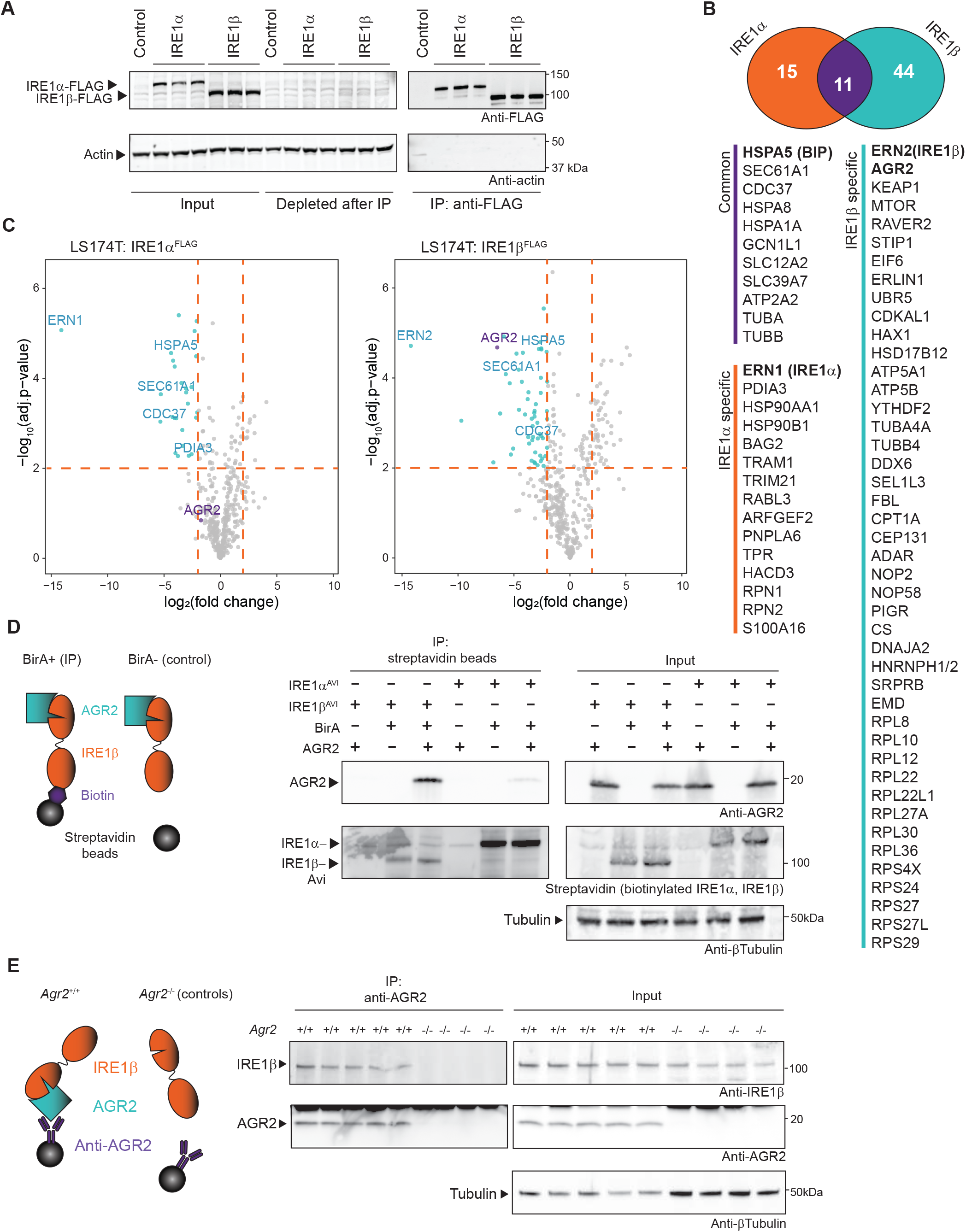
The mucin chaperone AGR2 is a goblet cell specific interactor of IRE1β. **A** Verification of transgene expression and successful immunoprecipitation of FLAG-tagged IRE1 in the samples analyzed by MS in B and C. Lysates were probed for IRE1-FLAG expression using anti-FLAG, and actin was used as a loading control. **B** Proteins with a log_2_FC enrichment of >2 and log_10_Adj p-val of >2 for IRE1α-FLAG and IRE1β-FLAG immunoprecipitation (IP) compared to control cells. The Venn diagram shows the number of proteins that were detected uniquely associated with one of the two IRE1 paralogues or that were commonly identified with both IRE1 paralogues. **C** Volcano plot depicting the cutoff criteria and significantly enriched proteins in each IP. X-axis shows the log_2_ fold change of the measured peptide intensities of a given protein in the control condition over the IRE1 IP condition. Y-axis shows the FDR corrected p-value obtained by two sample t-test in Perseus. **D** Confirmation of specific interaction between AGR2 and IRE1β, but not IRE1α. IRE1 proteins were tagged with an Avi-tag that is specifically biotinylated upon BirA co-expression. The biotinylated Avi-tag was precipitated using streptavidin beads. For control conditions, BirA was omitted. Blots were probed for co-precipitation of AGR2 and streptavidin to detect Avi-tag biotinylated IRE1. Tubulin was used as a loading control. Representative of 2 independent experiments. **E** Confirmation of the AGR2-IRE1β interaction in murine tissue. Colons were isolated and digested, and IP was performed using anti-AGR2. *Agr2* deficient mice were used as a negative control to assess whether IRE1β binds aspecifically to the antibody/bead complex. IP samples were probed for IRE1β co-precipitation via immunoblot. Tubulin was used as a loading control.

### Co-expression of AGR2 dampens endonuclease activity of IRE1β

Next, we assessed the endogenous expression levels of AGR2 in the different cell lines we previously analyzed for IRE1β expression (**Fig 1A** and **Fig S2A**). This revealed that LS174T cells show a prominent expression of AGR2, while Calu-1 cells do not (**Fig S2A**). HeLa cells, which die upon overexpression of IRE1β ^28^, also do not express AGR2 endogenously (**Fig S2A**). This suggests that AGR2 could be a modulator of IRE1β signaling in goblet cells by providing protection from unrestrained endonuclease activity and toxicity. Along these lines, other members of the PDI family have been implicated in regulating ER stress responses ^33–36^. To test this hypothesis, we monitored endonuclease activity-dependent cell death in Calu-1 cells upon co-expression of IRE1β and AGR2. Calu-1^ERN^^1-/-IRE1βFLAG-DOX^ cells were transduced with a constitutively expressing AGR2 lentiviral construct (‘Calu-1^AGR2^’) or ER-targeted BirA as control protein (‘Calu-1^Control^’). Endonuclease activity, as measured by XBP1 splicing, was severely attenuated in Calu-1^AGR^^2^ cells compared to Calu-1^Control^ cells (**Fig 3A** and **Fig S2B**), while the effects on the canonical RIDD target *Bloc1s1* appeared less pronounced (**Fig 3A**). Of note, co-expression with AGR2 almost completely reversed the cellular toxicity triggered by IRE1β expression (**Fig 3B**) as quantified by Annexin V/Live-Dead staining via flow cytometry in cells either expressing or lacking AGR2 (**Fig 3C** and **Fig S2C**). After doxycycline-induced expression of IRE1β, protein levels in surviving cells were analyzed by SDS-PAGE (**Fig 3D**). This revealed that any remaining Calu-1 cells showed very low IRE1β expression levels (lane 2), unless 4µ8C was added to the cultures to block IRE1β endonuclease activity (lane 3), in line with the notion that none of the IRE1β-expressing cells survived in the culture dish when IRE1β activity was not inhibited. Upon co-expression of AGR2, IRE1β expressing cells could be readily retrieved (lanes 5-6), and inhibition of endonuclease activity by 4µ8C treatment only increased the amount of IRE1β that was retrieved marginally.

**Figure 3.**
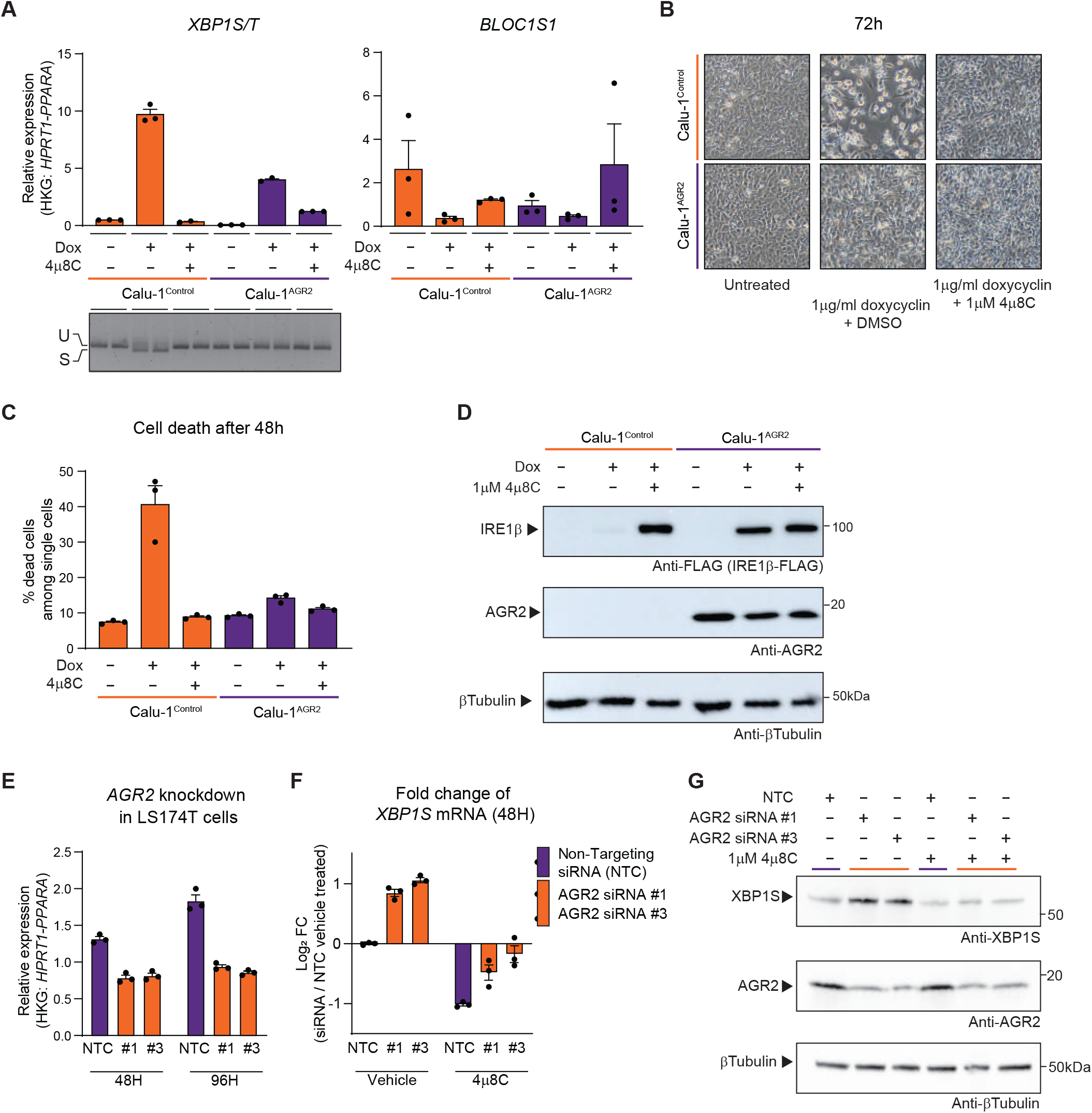
Co-expression of AGR2 dampens endonuclease activity of IRE1β. Calu-1*^ERN^*^1-/-IRE1βFLAG-DOX^ cells were transduced with a constitutive AGR2 transgene (“Calu-1^AGR^^2^“). Cells denoted as “Calu-1” are the original Calu-1*^ERN^*^1-/-IRE1βFLAG-DOX^ cells. **A** RT-qPCR analysis of *XBP1*S/T and *BLOC1S1* transcript levels after 24 hours of transgene induction using 1 μg/ml doxycycline. Bottom picture shows XBP1 splicing in the same samples assayed by conventional PCR. Representative of three independent experiments with three replicates per condition. **B** Photographs showing the phenotype of cultures overexpressing IRE1β-FLAG with and without exogenous expression of AGR2. Left panels show untreated cultures, middle panels show cultures that received 1μg/ml doxycycline for 72 hours, cultures in the right panels received 1μg/ml doxycycline and 1μM 4μ8C. Representative of three independent experiments. **C** Quantification of cell death in cultures overexpressing IRE1β-FLAG with and without exogenously added AGR2 after 48 hours. All cells in the culture dish were stained with AnnexinV and Live/Dead stain and analyzed by flow cytometry. All single and double positive cells were considered as dead cells. Representative of 2 independent experiments with three replicates per condition. **D** Analysis of IRE1β-FLAG and AGR2 expression in the cell lines used for C. Only cells that remained attached in the dish were collected and lysates were probed for IRE1β expression using anti-FLAG and AGR2 expression using anti-AGR2. Tubulin was used as a loading control. **E** Validation of AGR2 knockdown efficiency in LS174T^ERN^^1-/-IRE1βFLAG-DOX^ cells. NTC is a non-targeting control pool of siRNA’s, #1 and #3 are siRNA’s targeting AGR2. **F** Changes in XBP1 splicing after AGR2 partial knockdown and/or treatment with 4μ8C or DMSO (vehicle). Splicing is shown as a log_2_ fold change over the NTC/vehicle treated cells. **E** and **F** are representative of three independent experiments with three replicates per condition. **G** Western blot confirmation of (E) and (F). Proteins were extracted after 72 hours and probed for XBP1S, AGR2 and tubulin expression.

On the other hand, when endogenous AGR2 was (partially) depleted from LS174T cells via siRNA (**Fig 3E**), XBP1 splicing was elevated, which was apparent both at mRNA (**Fig 3F)** and at protein level **(Fig 3G)** and could be reversed by co-treatment with 4µ8C (**Fig 3F, G**). Of note, 4µ8C also inhibited the basal XBP1 splicing levels in LS174T cells in non-targeting siRNA control conditions, showing that even in presence of endogenous AGR2 levels, IRE1β retained some baseline activity (**Fig 3F**). In summary, these data establish AGR2 as a direct modulator of IRE1β endonuclease activity in goblet cells.

### AGR2 blocks IRE1β activity through disruption of IRE1β dimers

We next addressed how AGR2 mediates IRE1β inhibition. Previously, PDIA6 and PDIA1, two other members of the large family of PDI’s, have been implicated in reversing IRE1α oligomerization ^34,37^, and we hypothesized that AGR2 could operate in a similar manner. To investigate this possibility, gel filtration assays were set up in two different cell lines, HEK293T cells or Calu1^ERN^^1-/-^ cells (**Fig 4A**), in which the oligomerization status of overexpressed IRE1β could be monitored. The HEK293T cells were engineered to transiently overexpress an IRE1β construct fluorescently tagged with monomeric superfolder GFP (msfGFP). The Calu1^ERN^^1-/-^ cells were stably expressing a doxycycline-inducible IRE1β-FLAG protein (**Fig 4A**). IRE1β-msfGFP eluted in 2 peaks: a large peak at ∼14.2mL and a smaller peak at ∼15.3mL (**Fig 4B**, dotted lines). These two species shifted in intensity when AGR2 was co-expressed: the first peak shifted to approximately ∼14.5mL and became less abundant, while the peak at ∼15.3 mL became more prominent (**Fig 4B**, orange and purple traces). Based on the previously established elution profile of IRE1β in dodecylmaltoside ^21^, the later elution fraction most likely represented monomeric IRE1β-msfGFP, while the larger fraction most likely represented dimeric IRE1β-msfGFP, indicating that co-expression of AGR2 shifted the IRE1β equilibrium towards the monomeric forms. It should be noted that this experiment was performed on cell lysates, hence formation of oligomers or binding of the dimers to other cellular proteins could not be excluded. When cleared protein lysates of Calu-1^Control^ and Calu-1^AGR^^2^ cells were fractionated by gel filtration into 0.2ml fractions, similar findings were obtained (**Fig 4C**). The early eluting fraction remained present even upon AGR2 co-expression, but a late-eluting (most likely monomeric) fraction of IRE1β-FLAG could only be observed in Calu-1^AGR^^2^ cells, but not in Calu-1^Control^ cells. AGR2 preferentially co-eluted with these smaller molecular weight fractions, again indicating that the interaction of AGR2 with IRE1β appeared to disrupt IRE1β dimers and/or oligomers (**Fig 4C**). While gel filtration is generally feasible for assessing the oligomerization status of full length IRE1 proteins, we could not be fully confident that a protein complex between a membrane-based protein such as IRE1β and an ER luminal protein like AGR2 would not be affected by the gel filtration process, where a small sample is diluted into a larger column volume. Additionally, many other cellular proteins may bind IRE1β-msfGFP, making firm conclusions on the mono- or dimeric status challenging. We therefore explored an alternative setup to assess whether AGR2 could disrupt IRE1β dimers. To this end, a co-immunoprecipitation experiment was designed in which IRE1β was expressed fused to either the Avi tag or a FLAG tag (**Fig 4D**). Efficient co-precipitation of IRE1β-FLAG species with the biotinylated Avi-tagged IRE1β bait indicated the presence of at least IRE1β dimers. Co-expression of AGR2 led to a disruption of the IRE1β-dimers as observed by a reduction of FLAG-IRE1β signal co-precipitating with Avi-IRE1β in favor of AGR2 co-precipitation (**Fig 4E**, compare lanes 2 and 3). This indicated that AGR2 was able to shift IRE1β complexes that were at least dimeric in size to a monomeric state. This molecular transition was dose-dependent, with a >4 molar excess of AGR2 over FLAG-IRE1β having the strongest effect (**Fig 4F**). Collectively, these findings revealed that AGR2 expression disfavored the formation of IRE1β dimers.

**Figure 4.**
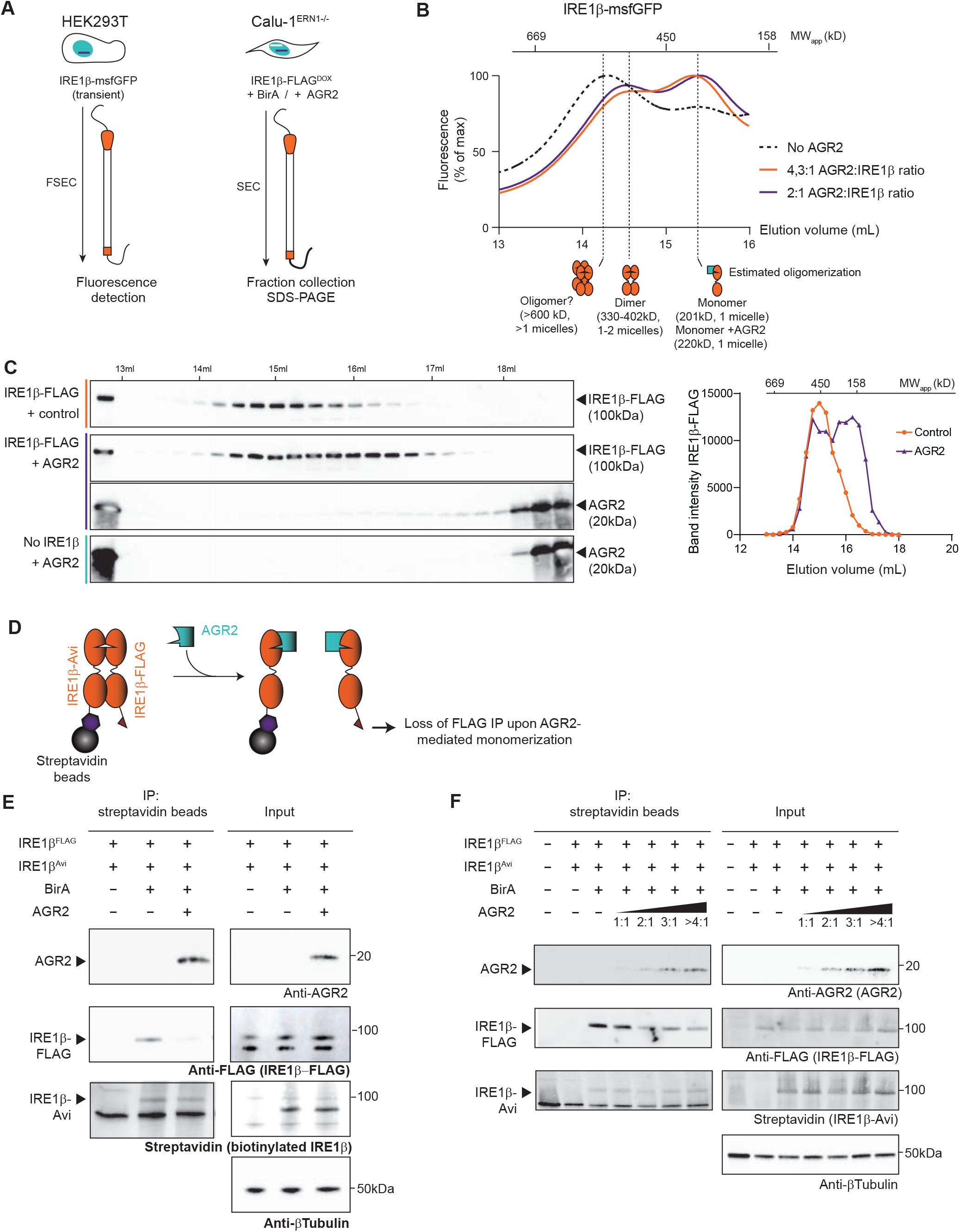
AGR2 blocks IRE1β activity through disruption of IRE1β dimers. **A** Schematic overview of gel filtration experiments in B and C. **B** msfGFP fluorescence measured during elution of HEK293T lysates overexpressing IRE1β in absence of AGR2 (black dotted trace) or in presence of AGR2 (orange and purple traces indicating different ratios of transfected AGR2:IRE1β plasmid). Top scale represents approximate elution profile and expected MW of protein standards. Bottom drawings indicate expected oligomerization status based on protein standards and the previously obtained elution profile (Grey). Experiment performed once. **C** IRE1β-FLAG expression in fractions collected after gel filtration of protein lysates from Calu-1^ERN^^1-/-IRE1βFLAG-DOX^ cells, in the absence or presence of additional AGR2 expression. Line graph shows quantification of band intensities from the gel. Representative of two independent experiments. **D** Schematic representation of competition IP experiments in E and F. IRE1β is expressed with an Avi-tag or FLAG tag in equimolar amounts. After biotinylation of the Avi-tag by BirA, both the Avi-tag and FLAG-tag will be detected after streptavidin IP if dimers have been formed. If addition of another protein (*e.g.* AGR2) would block this process, a loss of signal is expected. **E** Competition IP showing loss of dimer formation upon co-expression of AGR2. Samples were immunoblotted with anti-AGR2, anti-FLAG and Streptavidin. Tubulin was used as a loading control in input samples. Representative of two independent experiments. **F** Competition IP demonstrating concentration-dependent loss of dimer formation upon increasing AGR2 co-expression. Samples were immunoblotted with anti-AGR2, anti-FLAG and Streptavidin. Tubulin was used as a loading control in input samples. Representative of two independent experiments.

### The catalytic-dead C81S and disease-causing H117Y mutations in AGR2 abrogate its ability to bind and inhibit IRE1β activity

In literature, several key residues have been identified in AGR2 that impact its function and/or tertiary structure (**Fig 5A**). As AGR2 is characterized by an unusual CXXS thioredoxin catalytic motif, a single point mutation (C81S) is sufficient to disrupt its thiol exchange ^29^. The resolution of the nuclear magnetic resonance (NMR) structure of AGR2 revealed that AGR2 exists in a dimer, and that the single amino acid substitution mutations E60A and K64A behave as obligate monomers ^38^. Recently, the amino acid substitution H117Y has been identified as disease-causing in families where homozygous individuals display severe early-onset inflammatory bowel disease (IBD) ^30,39^. We wondered whether AGR2 requires dimerization and its thioredoxin motif to mediate the interaction with IRE1β. To this end, all the mutations were assayed in a similar co-IP setup as performed in **Fig 4F**, to assess simultaneously whether an AGR2 mutant retained binding to IRE1β and/or the ability to disrupt IRE1β dimers (**Fig 5B**). This revealed that the monomeric E60A and K64A mutants retained the capacity to bind IRE1β and consequently, to disrupt the formation of IRE1β-Avi/IRE1β-FLAG dimers. On the other hand, both the C81S and H117Y mutations lost interaction with IRE1β, which was reflected by a sustained formation of IRE1β-Avi/IRE1β-FLAG dimers, similar as in conditions without exogenous added AGR2 (**Fig 5C**).

**Figure 5.**
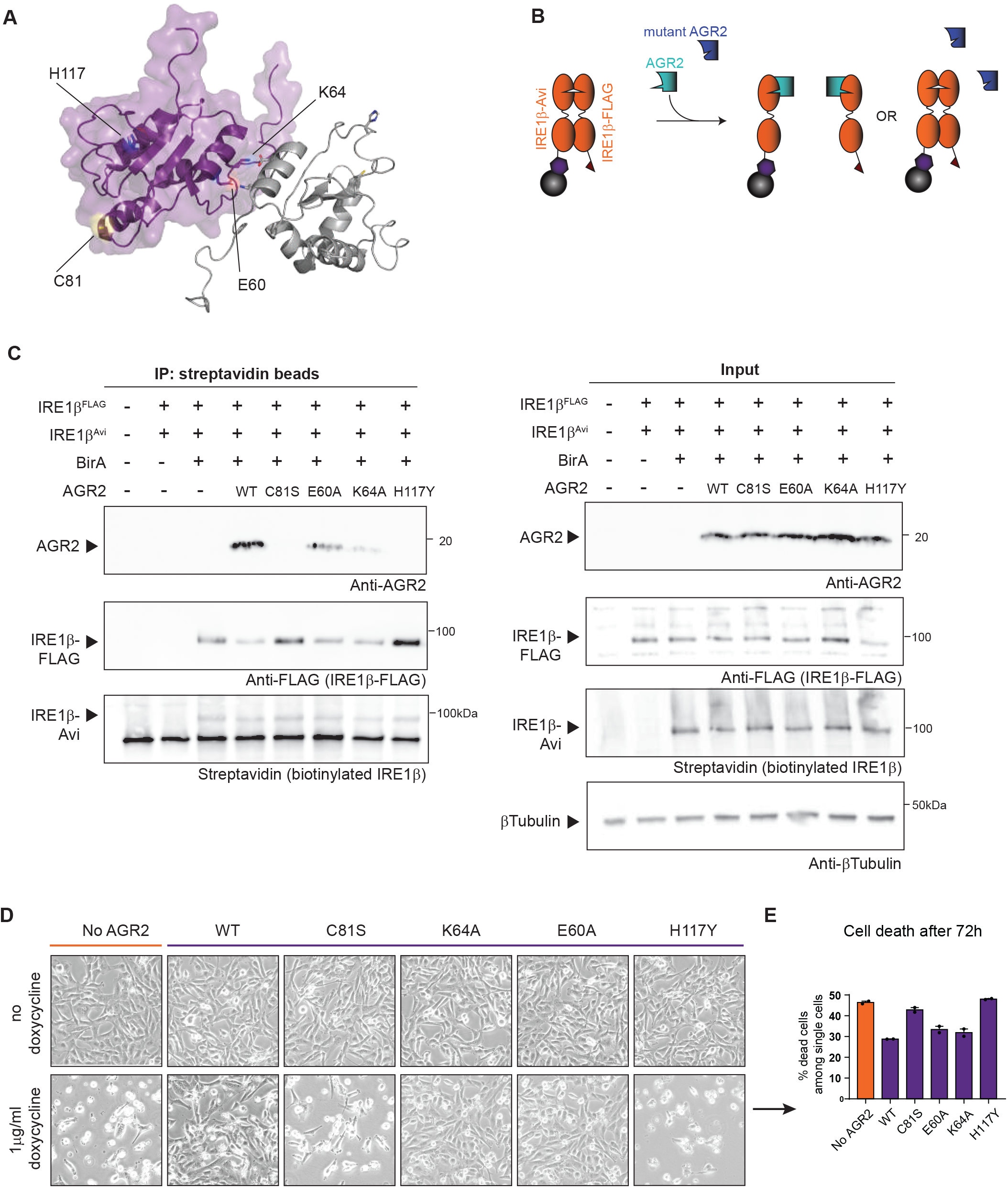
The catalytic-dead C81S and disease-causing H117Y mutations in AGR2 abrogate its ability to bind and inhibit IRE1β activity. **A** The structure of AGR2 (pdb: 2LNS) visualized in PyMol with the relevant mutations indicated. Purple and grey cartoons depict two AGR2 molecules and their dimer structure ^38^, specific residues are represented as ball-and-sticks. **B** Schematic overview of competition IP using AGR2 mutants. **C** Competition IP showing loss of dimer inhibition using C81S and H117Y AGR2 mutants. Samples were immunoblotted with anti-AGR2, anti-FLAG-IRE1β and Streptavidin. Tubulin was used as a loading control in input samples. Representative of two independent experiments. **D** Calu-1^ERN^^1-/-IRE1βFLAG-DOX^ cells were transduced with a constitutive AGR2 transgene (wild-type or the indicated mutants) and IRE1β-FLAG overexpression was induced with 1μg/ml doxycycline for 72 hours. **E** Quantification of cell death in cell lines from D after 72 hours of transgene expression. All cells in the culture dish were stained with Annexin V and Live/Dead stain and analyzed by flow cytometry. All single and double positive cells were considered as dead cells. Representative of 2 independent experiments with two replicates.

Based on the observation that WT AGR2 attenuates IRE1β-mediated cytotoxicity, we speculated that AGR2 mutations that block binding of AGR2 to IRE1β would unleash IRE1β endonuclease activity. To assess this hypothesis, all AGR2 mutants were transduced as constitutive lentiviral constructs in the Calu-1^ERN^^1-/-IRE1βFLAG-DOX^ cell line. All lines show similar expression of AGR2 protein (**Fig S3A**). The capability of each mutant to disrupt the catalytically active IRE1β dimer (**Fig 5C**) correlated with their ability to inhibit IRE1β-mediated cell death (**Fig 5D, E**), indicating that the protection of AGR2 against IRE1β endonuclease-mediated cytotoxicity was strictly dependent on its ability to interact with IRE1β. Overall, our data establish AGR2 as a goblet cell specific regulator of IRE1β endonuclease activity by disrupting IRE1β dimers and shifting the protein towards a monomeric, catalytically inactive state. In cells that do not express AGR2, overexpression of IRE1β leads to unrestrained endonuclease activity and cell death.

## Discussion

Goblet cells, embedded in the epithelia lining the mucosal interfaces of the respiratory and gastrointestinal track, are specialized secretory cells involved in the production and secretion of the gel-forming mucins MUC5A/B and MUC2, the major constituents of mucus. The proper folding and assembly of the 5100 aa MUC2 proteins is a real “tour-de-force”, following a highly structured process involving the formation of multiple disulfide bridges at the C- and N-termini, dimerization and trimerization steps in the ER and Golgi respectively, and massive O-glycosylation of the central Proline, Threonine and Serine (PTS) domain ^2^. This specialized function is reflected by the goblet cell transcriptomic and proteomic profile, which highlights activation of cellular pathways associated with protein production, folding, vesicle transport, glycosylation and secretion ^40^. Some of these genes, such as *Clca1, Zg16, Fcgbp1, Ern2, Agr2* or the transcription factors *Atoh1* or *Spedf* appear uniquely expressed in goblet cells ^41^. Amongst them, the protein disulfide isomerase (PDI) AGR2 is one of the most abundantly expressed proteins in the goblet cell ER ^40^.

AGR2 is a highly conserved protein, originally identified in Xenopus, where it was shown to play an essential role in development ^42–44^. It has an atypical KTEL ER-retention motif that localizes the protein to the ER ^45^, however, it has been found to be secreted as well ^46,47^. Several observations suggested a role for AGR2 in mucus homeostasis. Polymorphisms in the *AGR2* gene have been associated with increased risk of Ulcerative colitis and Crohn’s disease ^48^ and *Agr2*-/- mice lack an inner mucus layer in their gastrointestinal tract ^29,46^. An earlier study suggested that AGR2 would use its isomerase activity to form mixed disulfide bonds with MUC2, in this way assisting in mucus folding ^29^. However, this could not be confirmed by others ^46^ and up to now it remains unclear how AGR2 contributes to mucus homeostasis. As mentioned, the protein has been retrieved as an integral component of the intestinal mucus layer, indicative of a potential extracellular function as well ^47^.

We here identified AGR2 as a specific regulator of IRE1β, a sensor of the unfolded protein response (UPR) that is uniquely expressed in goblet cells, and that plays an essential role in goblet cell development and mucus homeostasis ^8,24,25^. AGR2 interacts specifically with IRE1β, but not with its ubiquitously expressed paralogue IRE1α, both upon overexpression in cell lines and in vivo in mouse colon tissue. As shown by Neidhardt *et al.* in a parallel manuscript, this interaction happens in a direct manner. Binding of AGR2 to IRE1β dampens its endonuclease activity, leading to a reduction in XBP1s splicing activity and RIDD. Besides, AGR2 also protects cells from IRE1β-mediated cytotoxicity, a hitherto unexplained phenomenon observed when IRE1β is exogenously expressed in non-goblet cell lines ^28^. Co-expression with AGR2 rescues cells from IRE1β-mediated cell death, in a manner that strictly depends upon their interaction, as any AGR2 mutant that loses interaction with IRE1β also loses its capacity to regulate its endonuclease output. On the reverse, siRNA-mediated reduction in AGR2 expression in goblet cells leads to increased IRE1β activity and XBP1 splicing. At the mechanistic level, we could demonstrate that AGR2 tends to disrupt IRE1β dimers, shifting IRE1β from a dimeric/oligomeric, highly active state towards a monomeric, inactive state. This was also confirmed by using recombinant proteins (Neidhardt *et al*.) and is conceptually highly reminiscent of how the Hsp70 chaperone BIP regulates IRE1α activity.

Currently, two alternative mechanisms have been put forward as to how BIP would engage the IRE1α luminal domain (LD), either in an ATP-dependent manner via its substrate binding domain as a true chaperone-substrate interaction ^49^, either in an ATP/chaperone-independent manner via its nucleotide binding domain ^50^. In the first model, unfolded proteins would compete with IRE1 LD for binding to the BIP substrate binding domain (SBD) (Amin-Wetzel et al., 2017, Preissler & Ron, 2019). In the second model, binding of unfolded proteins to the SBD of BIP would allosterically trigger the release of BIP from IRE1^9,50,51^. Independent of the upstream mechanism, both models would predict that the dynamic binding of BIP to IRE1α would establish a pool of inactive monomeric IRE1, against its intrinsic propensity to form dimers ^9^. Based on emerging data, BIP does not function isolated, but in cooperation with other chaperones such as members of the protein J family ^49^. In addition to BIP, also members of the large protein disulfide isomerase (PDI) family have been shown to regulate the activity of UPR sensors. The ER luminal oxidoreductase PDIA6 binds to oxidized Cysteine 148 in the luminal domain of IRE1α, which forms a disulfide bridge with Cys332 during activation-induced oligomerization ^52^. Reduction of the disulfide bridge by PDIA6 promotes inactivation of IRE1α and loss of PDIA6 leads to prolonged IRE1 activity with deleterious effects in *C. elegans* ^34^. Of note, a later study in mammals observed the opposite and proposed that PDIA6 would sustain long-term activation of IRE1 by an as-yet undefined mechanism ^35^. Finally, also (phosphorylated) PDIA1 has been shown to attenuate IRE1α activity by binding its luminal domain, this time in a cysteine-independent manner ^37^.

Neither Cys148, nor Cys332 are conserved in IRE1β (**Fig S4A**), but IRE1α Cys109 is conserved in the IRE1β luminal domain at position Cys117 ^53^. A second cysteine is present in the IRE1β luminal domain (Cys204), but, based on predictive modeling, neither cysteine appears to be solvent accessible in IRE1β. Furthermore, also AGR2 functions differently from canonical PDIs in the sense that it harbors only one single thioredoxin CXXS motif. It has been postulated to form dimers, that could allow catalytic activity ^38^. We found that the interaction of AGR2 with IRE1β occurred independently of AGR2 dimerization but depended on the presence of an active catalytic cysteine, as reflected by a loss of AGR2^C81S^ capacity to repress IRE1β cytotoxic activity. Similar findings are presented in an accompanying manuscript by Neidhardt *et al*. While the interaction between IRE1β and AGR2^C81^ mutants was weakened (though not fully disrupted), gel filtration assays on recombinant proteins demonstrated a diminished potency of AGR2 cysteine mutants to disrupt IRE1β dimers, in line with our own findings. Still, we can reasonably assume that AGR2 does not engage IRE1β through a mixed disulfide bond. Consistent with this notion, AGR2 was identified as an interactor of IRE1β in an immuno-affinity screen performed under reducing conditions, in which mixed disulfides would not be detected. Neither was the formation of a mixed disulfide between IRE1β and AGR2 observed upon co-immunoprecipitation of the two recombinant proteins (Neidhardt *et al*, accompanying manuscript). This suggests that the catalytic cysteine would rather mediate an allosteric effect. To get more insight into how these mutations might affect the interaction between IRE1β and AGR2, we modeled their interaction using AlphaFold2-Multimer (**Fig S4B, C**). While these models have only predictive value, it was surprising to note that all 5 seed models created by AlphaFold2 docked AGR2 to the same flexible loop of IRE1β (amino acids Ala318 to Leu353) with the AGR2 C81 and the clinically relevant H117 motif ^30^ facing the interaction site with IRE1β. This could explain why both mutants lost interaction with IRE1β. On the other hand, the C81 moiety in AGR2 appeared to be spatially separated from the two cysteines present in the luminal domain of IRE1β, again suggesting an allosteric rather than a true thiol exchange role for the C81. Finally, the two known amino acids involved in AGR2 dimerization (E60, K64) were located at the opposite site of the interaction motif with IRE1β, consistent with our immunoprecipitation experiments. Notably, BIP has been shown to bind to IRE1α at the same loop position, being held by both the flexible loop and the tail region in IRE1α ^11^. This could suggest that both BIP and AGR2 might show optimized affinity for the different flexible loop regions of IRE1α and IRE1β respectively, explaining their selective interaction. Why would the goblet cell specific UPR sensor IRE1β be regulated by a unique chaperone that is specific to goblet cells? The beauty of the UPR system is how it adapts folding load to match folding capacity by ensuring several negative feedback loops in different cellular contexts. One of the most well-known examples is the mammalian UPR sensor PERK, which upon activation due to dissociation of BIP, temporarily halts cap-dependent protein synthesis and in this way prevents further import of unfolded proteins in the ER ^54–56^. More recently, an alternative system has been identified that leads to trimming of the available mRNA pool by the endonuclease activity of IRE1, a process termed RIDD ^17,18^. RIDD is conserved from yeast to men and has been found downstream of both IRE1α and IRE1β ^57^. Of note, the upstream triggers of RIDD remain poorly understood, as it has often been described in (artificial) conditions of XBP1 deficiency, and/or in conditions of prolonged stimulation with ER stress-inducing agents such as tunicamycin. Hence, its physiological role remains debated. Still, in many cell types RIDD has been shown to target cell type specific substrates, such as insulin in pancreatic β-cells ^58^, immunoglobulin in plasma cells ^59,60^ or components of the antigen presentation machinery in dendritic cells ^61^, making a cell type specific and physiological role for RIDD likely. By making use of IRE1β deficient mice, Tsuru *et al.* demonstrated that IRE1β specifically degrades *Muc2* mRNA in goblet cells by RIDD ^25^. How this was regulated remained at the time unclear. We now add AGR2 as a missing piece in this puzzle (**Fig 6**). As a goblet cell specific chaperone, AGR2 might be perfectly suited to detect mucus folding load in the ER and to relay this information to IRE1β through a highly specific interaction with its luminal domain. This is highly relevant, as so far, the upstream triggers of IRE1β remain largely unknown, although they appear distinct from classical UPR triggers such as tunicamycin or thapsigargin^8^. We speculate that in conditions of high mucus folding load, AGR2 might dissociate from IRE1β, unleashing its propensity to dimerize. This could trigger IRE1β-mediated RIDD activity, in this way adapting *Muc2* mRNA availability to the MUC2 folding capacity in the ER. Several questions remain though, especially regarding the specific function of AGR2 in mucus folding, and the regulation of the AGR2/IRE1β interaction.

**Figure 6.**
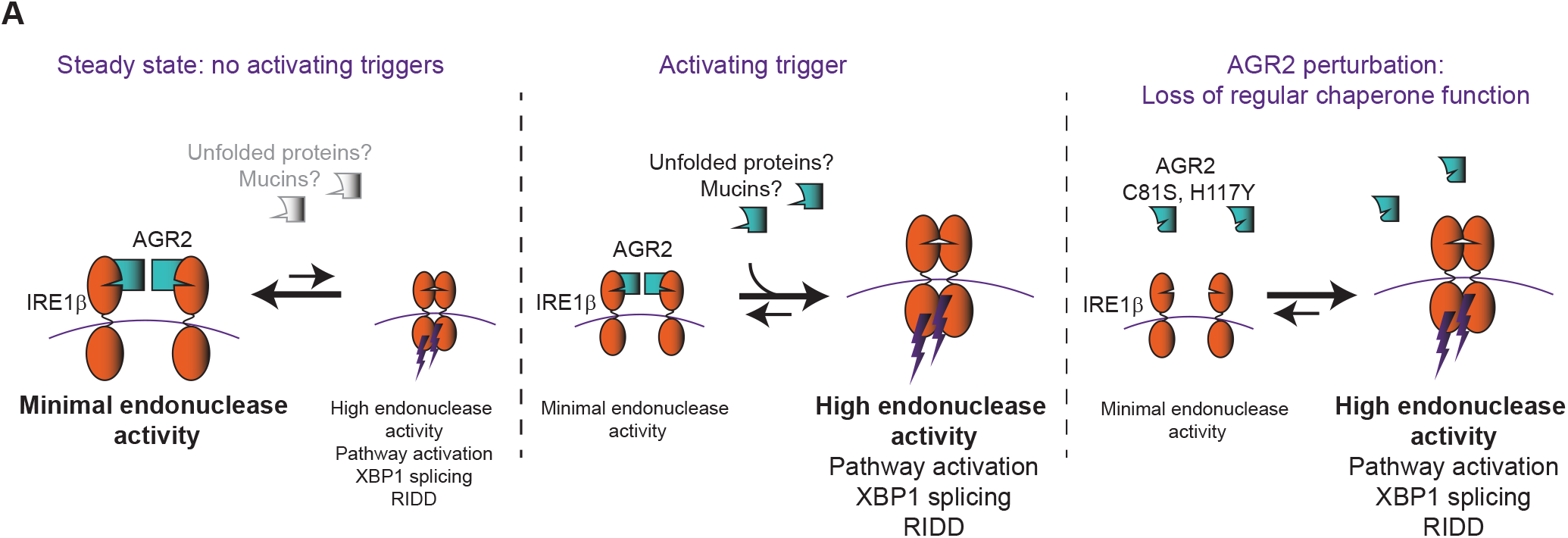
Proposed mechanism of IRE1β regulation by AGR2. In steady state conditions, AGR2 is bound to most IRE1β molecules and under these conditions, IRE1β is mainly present in the inactive, monomeric forms. As a result, overall IRE1β activity will be low. In conditions where an activating trigger is present (possibly unfolded MUC2 polypeptides, though this remains to be demonstrated), AGR2 is released from IRE1β in favor of binding other AGR2 chaperone substrates. As a result, IRE1β is released, activated and overall IRE1β activity will be high. In case of AGR2^C81S^ and AGR2^H117Y^, interaction with IRE1β is disrupted leading to spontaneous IRE1β dimerization triggering its activity.

Finally, *AGR2* has recently come into view as a disease-causing gene in a subset of patients displaying IBD-like symptoms, among others. A homozygous H117Y mutation was identified in twins with severe early-onset IBD ^30^, and several other families have been identified that display profound inflammatory phenotypes at their mucosal surfaces due to mutations in AGR2 ^39^. As we have demonstrated that AGR2^H117Y^ loses the ability to interact with and block IRE1β, it is reasonable to assume that these patients do not only exhibit disturbed AGR2 chaperone function but may also be characterized by IRE1β hyperactivity (**Fig 6**). Currently, it is unknown whether IRE1β hyperactivity causes symptoms in vivo and how this translates to patient care. Our findings highlight the importance of further understanding the AGR2/IRE1β axis in goblet cell physiology and disease.

## Figure legends Supplementary figures

**Figure S1.**
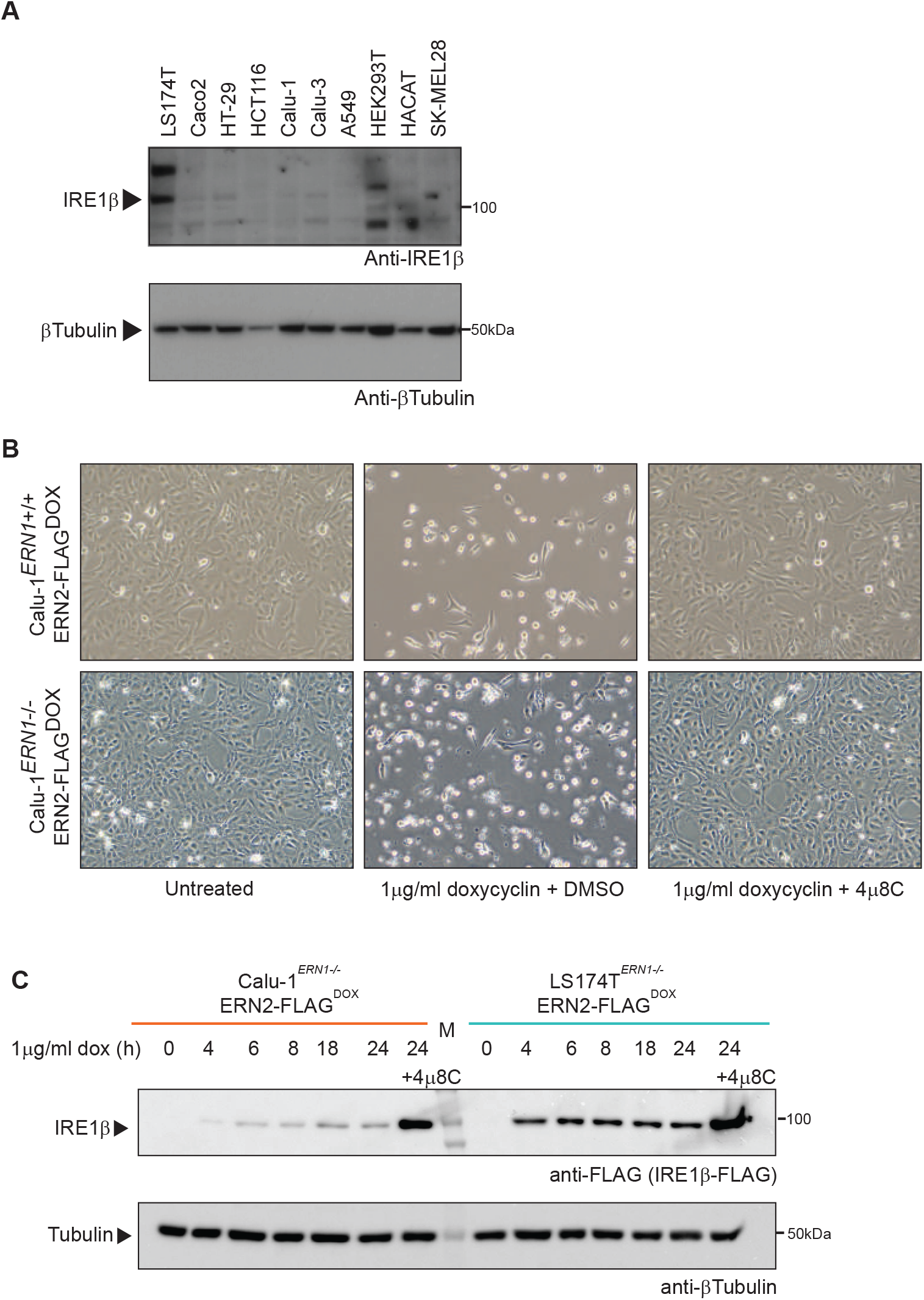
Validation of LS174T^ERN1-/-IRE1βFLAG-DOX^ and Calu-1 ^ERN1-/-IRE1βFLAG-DOX^ model systems. **A** IRE1β and AGR2 expression in cell lines. Proteins were extracted and probed for IRE1β expression and AGR2 expression via immunoblot. Tubulin was used as a loading control. **B** Photographs showing the phenotype of cultures overexpressing IRE1β-FLAG in IRE1α wild-type (*ERN1^+/+^*) and IRE1α deficient (*ERN1^-/-^*) cells. Left panels show untreated cultures, middle panels show cultures treated with 1μg/ml doxycycline for 72 hours, right panels show cultures treated with both 1μg/ml doxycycline and 1μM IRE1 endonuclease inhibitor 4μ8C. **C** Quantification of IRE1β transgene expression over time in Calu-1^ERN^^1-/-IRE1βFLAG-DOX^ and LS174T^ERN^^1-/-IRE1βFLAG-DOX^ cells by western blot. Cultures were treated with 1μg/ml doxycycline for the indicated times, protein lysates were probed for IRE1β-FLAG expression and tubulin as a loading control.

**Figure S2.**
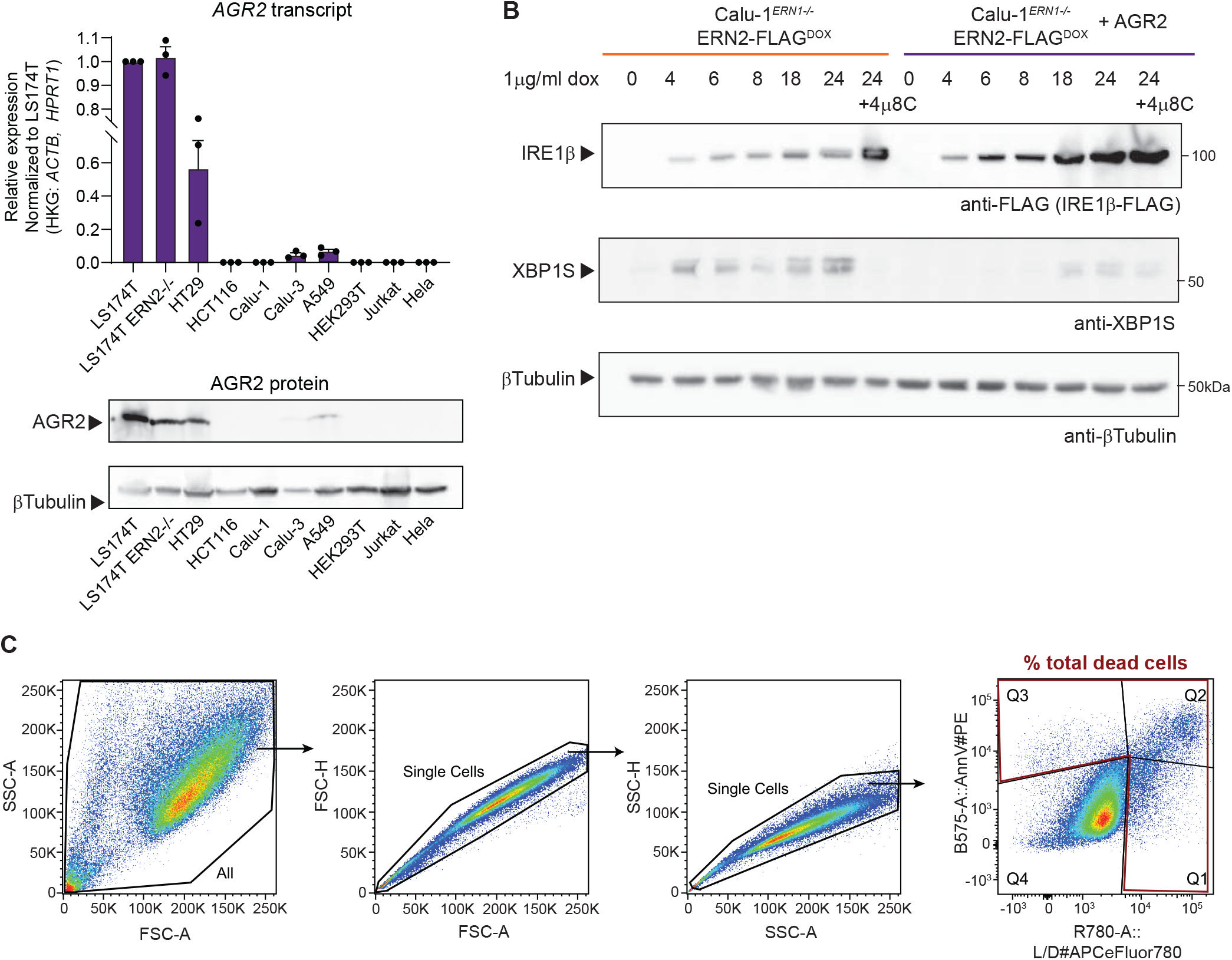
AGR2 expression is restricted to colon epithelial cell lines and affects Calu-1^ERN1-/-IRE1βFLAG-DOX^ phenotype. **A** *AGR2* transcript expression in cell lines assayed by RT-qPCR. Cultures were sampled at three different time points and *AGR2* expression is shown relative to the expression detected in LS174T parental cells. **B** Quantification of IRE1β-FLAG transgene expression over time by western blot in cell lysates derived from Calu-1^ERN^^1-/-IRE1βFLAG-DOX^ co-expressing ER-targeted BirA as a control protein (left, orange), or AGR2 (right, purple). Cells received 1μg/ml doxycycline to induce expression of IRE1β. Protein lysates were prepared at the indicated times and probed for IRE1β-FLAG expression, XBP1S and tubulin as a loading control. **C** Gating strategy to assess cell death. Doublets were gated out and dead cells were gated on via AnnexinV and Live/Dead positive staining. All cells staining positive for a single, or both cell death markers were considered dead (red gate).

**Figure S3.**
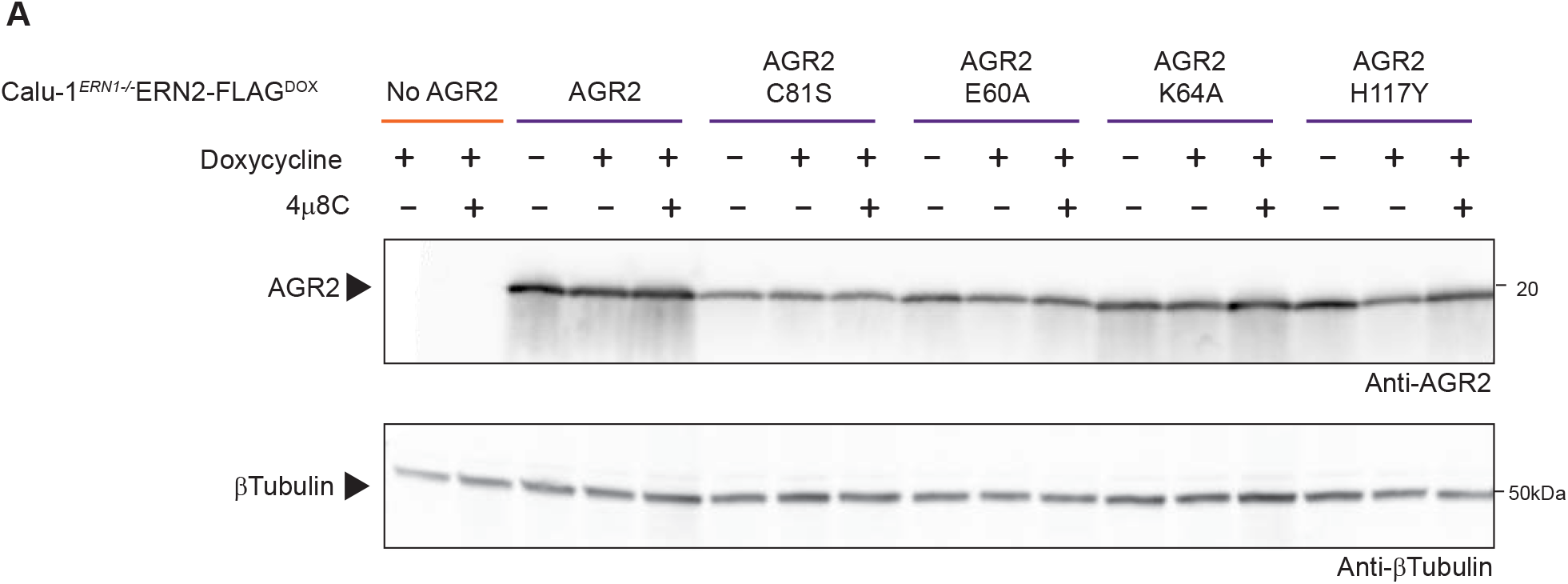
AGR2 expression in stably transduced cell lines. **A** AGR2 expression in cultures analyzed in Fig 5D and 5E. Tubulin was used as a loading control.

**Figure S4.**
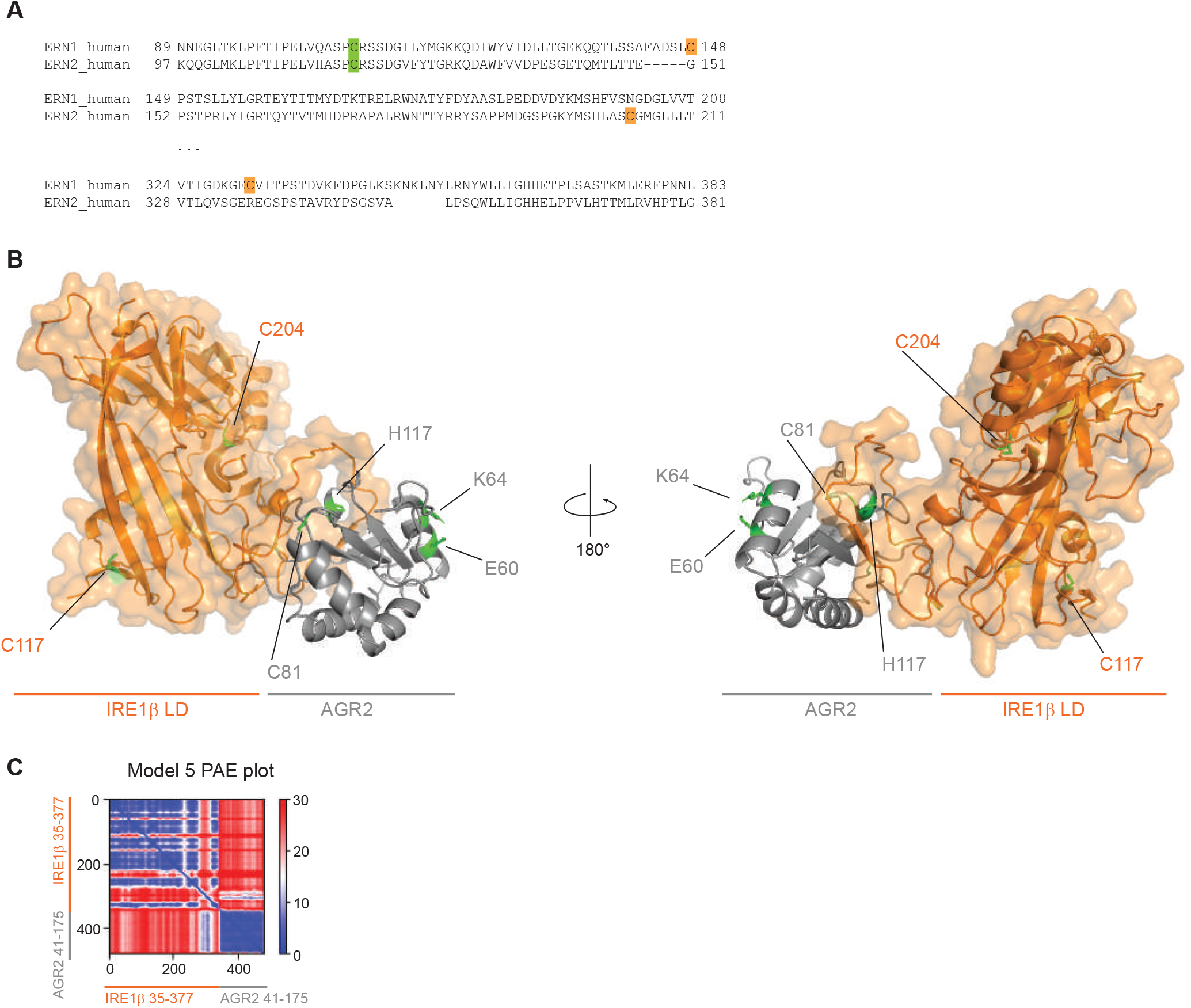
AGR2 is predicted to bind a flexible loop region in the IRE1β luminal domain. **A** BLAST alignment of the regions containing cysteines in human IRE1α and IRE1β. Green square indicates the sole conserved cysteine in IRE1α and IRE1β luminal domain, orange squares show cysteines present in only one of the paralogues. **B** Highest scoring AlphaFold2-Multimer model (pTM score = 0.662), modeled using IRE1β residues 35-377 (Uniprot Q76MJ5) and AGR2 residues 41-175 (Uniprot O95994). The IRE1β luminal domain is shown in orange and AGR2 in grey. Labels indicate the highlighted green residues. **C** Predicted aligned error (PAE) plot for the model shown in B.

## Methods

### Cell culture and compounds

Calu-1, HCT116 and HT29 cells were cultured in modified McCoys 5A (Gibco) supplemented with 10% FBS. LS174T and Calu-3 cells were cultured in Minimum Essential Media (MEM) (Sigma) and supplemented with 1XGlutamax (Gibco), 1XNEAA (Gibco), 1XNa-pyruvate (Gibco) and 10% FBS. HEK293T, A549 and HELA were cultured in Dulbecco’s Modified Eagle Medium (DMEM) (Gibco) and supplemented with 10% FBS. Jurkat cells were cultured in Roswell Park Memorial Institute (RPMI) (Gibco), supplemented with 10% FBS. All cells were obtained through ATCC or ECACC. All cells were cultured in a humidified 37 degree C incubator at 5% CO_2_.

To induce transgene expression in transduced cell lines (see below), cells were treated with 500ng-1µg/ml doxycycline (Sigma-Aldrich) in the culture media for the indicated times. 4µ8C (Sigma-Aldrich) was added at 1-5 µM to inhibit endonuclease activity where indicated.

### CRISPR/Cas9

Calu-1*^ERN^*^1^*^-/-^*cells were constructed by transfection with pSpCas9n(BB)-2A-Puro (Feng Zhang lab, Addgene #48139) containing sequences for guides targeting the start codon (WGE ID’s 1150311103 and 1150311093). After puromycin selection, cells were seeded as single cell subclones and clones with no detectable IRE1α protein were treated with tunicamycin to further verify lack of XBP1 splicing in response to tunicamycin. LS174T *^ERN^*^1^*^-/-^* cells and LS174T *^ERN^*^2^*^-/-^* cells were constructed by transfection with pSpCas9(BB)-2A-GFP (Feng Zhang lab, Addgene #48138) containing sequences for guides targeting exon 8 (WGE ID’s 1150303698 and 1150303744 for ERN1, and ID’s 1135590972 and 1135590916 for ERN2). After puromycin selection, cells were seeded as single cell subclones and clones with no detectable IRE1α protein were treated with tunicamycin to further verify lack of XBP1 splicing in response to tunicamycin. For assessing IRE1β expression, clones were genotyped with primers (Fwd: 5’-GAGTTGCTGAACAGTGGGGG-3’, 5’-GTGTAGACGCCCATCACAGG-3’, Rev: 5’-GACCTTACAGGTGCCCCAAG-3’) to verify disruption of exon 8 and these clones were further verified by western blot.

### Plasmids

IRE1β-FLAG was ordered through gene synthesis (Gen9), and IRE1α-FLAG was a kind gift from Sarah Gerlo (Ghent University). IRE1-FLAG sequences were amplified with AttB sites, cloned into pDONR223 using BP clonase II (Invitrogen) and transferred into pDEST51 (Invitrogen) using LR clonase II (Invitrogen). For BirA-Avi co-IP experiments, full-length IRE1α and IRE1β were PCR-amplified with a C-terminal KpnI-SpeI site, flanked by AttB sequences and transferred to pDONR221 with BP clonase II (Invitrogen). The Avi tag fragment was obtained through annealing of two oligo’s (Integrated DNA Technologies) with the correct overhangs (Top strand: 5’-CGGCGGTGGCCTGAACGACATCTTCGAGGCTCAGAAAATCGAATGGCACGAAA-3’, bottom strand: 5’-CTAGTTTCGTGCCATTCGATTTTCTGAGCCTCGAAGATGTCGTTCAGGCCACCGCCGGTAC-3’) and cloned into the KpnI-SpeI site. The sequences were transferred to pDEST51 or the inducible Virapower TRex expression system (Invitrogen) with LR clonase II (Invitrogen). IRE1β-msfGFP was produced by PCR ampification of IRE1β with a C-terminal KpnI-SpeI site. The msfGFP sequence ^62^ was ordered through gene synthesis (IDT) and PCR-amplified with KpnI-SpeI cloning sites and cloned into the IRE1β KpnI-SpeI sites. AGR2 was PCR amplified from LS174T cDNA with AttB sites and transferred in pDONR221 and pLenti-DEST-EF1A-Hygro. All indicated mutations were introduced with the QuickChange protocol (Agilent) and sequences were verified by Sanger sequencing. BirA was PCR amplified from pDisplay-BirA-ER (Alice Ting lab, Addgene #20856) with AttB sites as described above and transferred to pDEST12.2 or pLenti-DEST-EF1A-Hygro. Plasmid preparations were performed using Nucleobond Midi Xtra (Machery-Nagel).

### Viral vectors and transduction

Lentiviral particles were prepared in HEK293T cells using standard calcium-phosphate transfection of the lentivector with packaging plasmids pMD2.G (Trono lab, Addgene #12259) and pCMV-dR8.74 (Trono lab, Addgene #22036). Viral particles were concentrated through ultracentrifugation. Cell lines were transformed with both the pLenti3.3 repressor at MOI5, the IRE1β-FLAG transgene in pLenti6.3/TO/DEST at MOI2 and constitutive AGR2 at MOI2 or greater. Calu-1 cells carrying stable integration of the transgene were selected using G418 (0.5mg/ml), blasticidin (10ug/ml) and hygromycin (200ug/ml) respectively. LS174T cells were selected using G418 (0.4mg/ml), blasticidin (8ug/ml)

### AP-MS

LS174T cells were plates in 2x 145cm^2^ culture dishes/repeat pulldown (a total of 3 repeats was carried out per conditions). Transgene expression was induced using 1µg/ml doxycycline. After 24H, cells were collected in PBS. Cell pellets were washed in PBS, weighed and lysed in 400µl/100µg cell pellet AP-MS buffer (50mM HEPES-KOH pH8.0, 100mM KCl, 2mM EDTA, 0.1% NP40, 10% glycerol, 1mM DTT, 0.5mM PMSF, 0.25mM sodium orthovanadate, 50mM β-glycerophosphate, 10mM NaF and protease inhibitor cocktail (Roche)). Cells were subjected to one freeze-thaw cycle after which lysates were cleared by centrifugation (16 000g, 4DC, 20’) and transferred to a new tube where they received BioM2 anti-FLAG (Sigma-Aldrich) loaded Dynabeads MyOne Streptavidin T1 (Invitrogen) (5µl anti-FLAG with 50µl beads per combined set of 2x 145cm^2^ dish) and were incubated with end-over-end rotation at 4DC for two hours. Beads were washed, peptides were eluted by on-bead trypsin (Promega) digest and acidified for MS analysis. 2.5µl sample was injected on a Q Exactive hybrid Quadrupole-Orbitrap analyzer (ThermoScientific). Protein identification was carried out using the Andromeda search engine of the MaxQuant v1.6.3.4. software package ^63^ and the SwissProt human proteome (versions 08-2018). Identified proteins were quantified using label-free quantification and significant enrichment of proteins was determined after imputation of missing values using a Student’s paired t-test in Perseus v1.5.5.3. Proteins were considered to be enriched when they were characterized by a log_2_ fold change (log_2_FC) enrichment of >2, and log_10_Adj p-val of >2.

### Co-immunoprecipitation (co-IP)

HEK293T cells were seeded in 6 well plates at 0,5.10^6^ cells/well and transfected the next day by calcium phosphate transfection (6.6ug DNA/transfection) in culture medium containing 1μM biotin. Unless stated otherwise, cells were transfected with a ∼2.5:1 molar ratio of AGR2:IRE1 plasmid. To mediate biotinylation of the IRE1α or IRE1β-Avi tag, BirA was included the transfection mix, and replaced by pSV-sport empty vector for control conditions. For competition coIP, IRE1β-Avi and IRE1β-FLAG plasmids were included in equal molar amounts. Cells were lysed in lysis buffer (1% NP-40, 10% glycerol, 250mM NaCl, 20mM HEPES pH7.9, 1mM EDTA with cOmplete mini protease inhibitor cocktail and PhosSTOP (Roche)). Cleared lysates were incubated for 1 hour at 4DC with Dynabeads Streptavidin T1 (Invitrogen) (8µl beads per sample). Beads were washed and proteins were eluted directly in loading buffer by incubating beads in loading buffer at 65DC. SDS-PAGE was performed, proteins were transferred to nitrocellulose membrane (Cytiva) and revealed using following antibodies: rabbit AGR2 (Cell Signaling Technologies, Clone D9V2F, 1/1000), mouse FLAG-M2-HRP (Sigma, A8592, 1/1000), Streptavidin-HRP (CST, 3999S, 1/1000), rabbit β-Tubulin-HRP (Abcam, ab21058, 1/5000), and anti-rabbit-HRP rabbit IgG-HRP (Dako, P0448, 1/2000). Images were captured on an Amersham 600 Imager (GE Healthcare).

### In vivo coIP

AGR2^tm1.2^Erle/J mice were purchased from the Jackson Laboratory. All mice were housed in SPF conditions under the current EU animal housing guidelines. Mice were euthanized by CO2 asphyxiation, and colons were extracted. After two washing steps in PBS to remove fecal matter, colon pieces were transferred to 1xHBSS containing 0.15% DTT to remove mucus. Pieces were then transferred to digestion buffer (RPMI supplemented with DNAse I and Liberase TM), homogenized with GentleMACS C tubes and incubated in a shaking warm water bath for 20 min. At 10 and 20 minutes, samples were again homogenized with GentleMACS C tubes. Single cell pellets were lysed in E1A buffer (1% NP-40, 10% glycerol, 250mM NaCl, 20mM HEPES pH7.9, 1mM EDTA with cOmplete mini protease inhibitor cocktail (Roche) and PhosSTOP (Roche)) and cleared by centrifugation. Anti-AGR2 antibody (clone 6C5, Santa Cruz Biotechnology) was first bound to Dynabeads Protein G (Invitrogen) according to the manufacturer’s instructions. Antibody-bead complexes were added to cleared lysates and incubated with rotation for 2 hours. Beads were washed and proteins were eluted directly in loading buffer by incubating beads in loading buffer at 65DC. SDS-PAGE was performed with Mini-Protean TGX 4-20% gels (Bio-Rad), proteins were transferred to PVDF membrane (Amersham 0.2 Hybond P, Cytiva) and revealed using following antibodies: mouse AGR2 (Santa Cruz Biotechnology, Clone 6C5, 1/1000), rabbit IRE1β (gift from David Ron, University of Cambridge, 1/2000), rabbit β-Tubulin-HRP (Abcam, ab21058, 1/5000), mouse IgG-HRP (Dako, P0447, 1/1500) and rabbit IgG-HRP (Dako, P0448, 1/2000). Images were captured on an Amersham 600 Imager (GE Healthcare).

### RT-qPCR

Total RNA was isolated from cells using the Aurum Total RNA Mini kit (Bio-Rad) according to the manufacturer’s instructions. cDNA was synthesized using SensiFast cDNA synthesis kit (Meridian Bioscience). RT-qPCR was carried out in 384-well plates using SensiFAST SYBR No-ROX kit (Meridian Bioscience) according to the manufacturer’s protocol. Preparation of mixes and transferring to 384 well plates was performed using the Janus automatic liquid sample handling station (Perkin-Elmer) or the I.DOT dispenser (Dispendix). All data analysis was performed by qbase+ (CellCarta). For each experiment, the most stable reference genes were selected using qbase+ and are mentioned in each figure or figure legend. Following primer pairs were used for RT-qPCR:

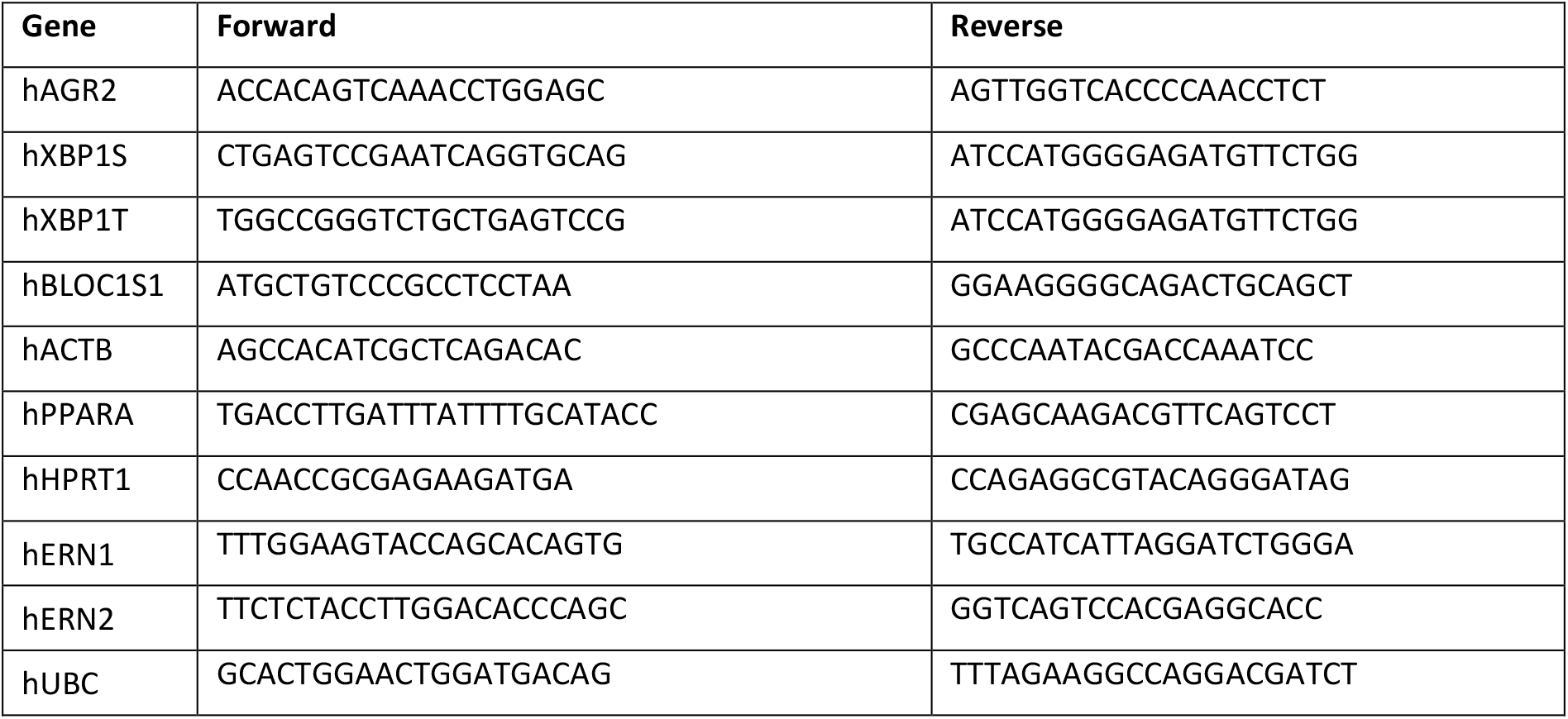

### XBP1 splicing assay

Total RNA was isolated and transcribed to cDNA as described for “RT-qPCR”. The XBP1 spliced region was amplified using Q5 high fidelity polymerase (NEB) and primers: fwd 5’-TGAAAAACAGAGTAGCAGCTCAGA-3’, rev 5’-TCTGGGTAGACCTCTGGGAGCTCC-3’. PCR reaction was separated on 2.5% agarose gel.

### Quantification of cell death

For flow cytometry, cells were seeded in 6 well plates at ∼80% confluency. The next day, IRE1β expression was induced using 1µg/ml doxycycline, supplemented with either 1µM 4µ8C or DMSO. 48h-72h later supernatant was collected, cells were washed in PBS which was added to the supernatant, and cells were dissociated with Accutase (Biolegend) and added to the supernatant. Cells were spun down and stained 20’ at 4DC with eBioscience™ Fixable Viability Dye eFluor™ 780 (Invitrogen), washed and stained 15’ at RT with Annexin V PE (BD Pharmingen^TM^) in annexin binding buffer. Cells were diluted in binding buffer and measured on a BD LSRII. Flow cytometry data was processed using FlowJo X.

### Gel filtration oligomerization assays

For Calu-1 cells, IRE1β expression was induced with 1μg/ml doxycycline for 16 hours. A 145cm^2^ culture dish at 80% confluency was lysed in 600ul lysis buffer buffer (150mM NaCl, 25mM Tris pH 8.0, 20mM dodecylmaltoside, 5mM β-mercaptoethanol with cOmplete mini protease inhibitor cocktail (Roche) and PhosSTOP (Roche)) at 4°C for 1 hour under continuous agitation. For assays in HEK293T cells, cells were seeded at 5.10^6^ cells/60cm^2^ dish and transfected with pDEST51-IRE1β-msfGFP plasmid alone or a 4.3:1 and 2:1 molar ratio of pDEST51-AGR2 and pDEST51-IRE1β-msfGFP plasmid (6.6ug in total) via Calcium-Phosphate transfection. Each dish was collected and lysed identically to the Calu-1 cells above. Lysates were cleared by centrifugation at 4°C, 16.000g for 20min. Fractionation of lysates was carried out on a Superose6 Increase 10/300 GL (Cytiva) size exclusion chromatography column equilibrated in running buffer (150mM NaCl, 25mM Tris pH 8.0, 0.5mM dodecylmaltoside, 5mM β-mercaptoethanol) at 0.5ml/min. Detection of fluorescent IRE1β-msfGFP was carried out on an RX-20Axs fluorescence detector (Shimazu, Ex: 485nm/Em: 510nm) in line with the UV detector. For IRE1β-FLAG detection, 0.25ml fractions were collected and assayed by SDS-PAGE (Criterion TGX 4-20% (Bio-Rad)) and western blot using HRP-coupled anti-FLAG M2 (Sigma, A8592) and anti-AGR2 (Cell Signaling Technologies, Clone D9V2F, 1/1000).

### Gene silencing

AGR2 knockdown in LS174T cells was obtained by reverse transfection of siRNA (siGENOME, Horizon Discovery, D-003626-01; D-003626-03; D-001206-14) using DharmaFECT1 transfection reagent (Horizon Discovery) according to manufacturer’s protocol. After 24H, culture medium was replaced and supplemented with 1µM 4µ8C or DMSO. 48H after transfection, cells were harvested. Total RNA was isolated, transcribed to cDNA and RT-qPCR was performed as described for “RT-qPCR” above. To assay AGR2 protein levels, cells were resuspended in lysis buffer (1% NP-40, 10% glycerol, 250mM NaCl, 20mM HEPES pH7.9, 1mM EDTA, 2mM DTT with cOmplete mini protease inhibitor cocktail (Roche) and PhosSTOP (Roche)) at 4°C for 15 minutes. Lysates were cleared by centrifugation at 4°C, 16.000g for 20min. Total protein was measured using Bradford assay (Bio-Rad). SDS-PAGE was performed on Mini-Protean TGX 4-20% gels (Bio-Rad). Proteins were blotted on Nitrocellulose membrane (Cytiva) and revealed using following antibodies: mouse anti-AGR2 (Santa Cruz Biotechnology, Clone 6C5, 1/1000, rabbit anti-XBP1s (Cell Signaling Technologies, clone E9V3E, 1/1000), rabbit β-Tubulin-HRP (Abcam, ab21058, 1/5000), anti-rabbit IgG-HRP (Dako, P0448, 1/2000), anti-mouse IgG-HRP (Dako, P0447, 1/1500). Images were captured on an Amersham 600 Imager (GE Healthcare).

### Structural modeling

All structure prediction models were generated using the Alphafold2.3.1 software ^64^ on an HPC cluster (Vlaams Supercomputer Centrum, Belgium). For each job 5 different seed models were requested that each were used to generate 5 models, giving rise to a total of 25 different models. All predictions were scored and ranked based on their PTM values. Figures were visualized using PyMOL ^65^.

## Acknowledgements and funding

SJ, SE and SNS are supported by the Vlaams Instituut voor Biotechnologie (VIB). This research was further supported by FWO (1228923N to EC, G017521N to SJ, and G049820N, G0H1222N, S000722N, S002322N to SNS), an ERC Consolidator grant (DCRIDDLE-819314 to SJ) and the Geconcerteerde Onderzoeksacties, Special Research fund, Ghent University (BOF16/GOA/023 to SE). LN is supported by Medical Research Council DTP and Gates Cambridge PhD programme. We would like to thank the VIB Proteomics core, the VIB Flow core and the VIB-IRC Tissue Culture Core for training, support and access to the instrument park and facility. We thank David Ron for feedback on the manuscript, and for providing IRE1β antiserum. We would like to thank Michael J. Grey for constructive discussions and advice.

## Author contributions

EC, SJ and SE conceived the study and designed experiments. EC and PG performed experiments with technical support from FF, EVDV and DDS. MP and SNS performed structural modeling and provided access to critical instrumentation. Manuscript was written by EC and SJ with significant input from SE, SNS, MP, PG and LN.

## Disclosure statement

The authors declare no competing interests.

## Notes

### Competing Interest Statement

The authors have declared no competing interest.

